# Experience-dependent coding of frequency-modulated trajectories by offsets in auditory cortex

**DOI:** 10.1101/613448

**Authors:** Kelly K Chong, Alex G Dunlap, Dorottya B Kacsoh, Robert C Liu

**Affiliations:** Wallace H. Coulter Department of Biomedical Engineering, Georgia Institute of Technology and Emory University, Atlanta, Georgia 30332, USA; Department of Biology, Emory University, Atlanta, Georgia 30322, USA; Center for Translational Social Neuroscience, Emory University, Atlanta, Georgia, 30322, USA

**Keywords:** pitch, belt, secondary auditory cortex, frequency modulation, neural plasticity, maternal behavior, neural coding, mouse, offset response, vocalization, USV, speech, music

## Abstract

Frequency modulations are an inherent feature of many behaviorally relevant sounds, including vocalizations and music. Changing trajectories in a sound’s frequency often conveys meaningful information, which can be used to differentiate sound categories, as in the case of intonations in tonal languages. However, it is not clear what features of the neural responses in what parts of the auditory cortical pathway might be more important for conveying information about behaviorally relevant frequency modulations, and how these responses change with experience. Here we uncover tuning to subtle variations in frequency trajectories in mouse auditory cortex. Surprisingly, we found that auditory cortical responses could be modulated by variations in a pure tone trajectory as small as 1/24th of an octave. Offset spiking accounted for a significant portion of tuned responses to subtle frequency modulation. Offset responses that were present in the adult A2, but not those in Core auditory cortex, were plastic in a way that enhanced the representation of an acquired behaviorally relevant sound category, which we illustrate with the maternal mouse paradigm for natural communication sound learning. By using this ethologically inspired sound-feature tuning paradigm to drive auditory responses in higher-order neurons, our results demonstrate that auditory cortex can track much finer frequency modulations than previously appreciated, which allows A2 offset responses in particular to attune to the pitch trajectories that distinguish behaviorally relevant, natural sound categories.

## INTRODUCTION

Frequency trajectory conveys meaning across many sounds, from environmental noises to vocalizations and music. In human speech, pitch modulation conveys expressive and linguistic meaning, especially in tonal languages [1-4]. In music, emotional meaning can be conveyed through pitch modulation [5]. In many species, such as birds, rodents, monkeys, dogs, and even some marine animals, the temporal features of a vocalization, including its pitch trajectory, vary by context, and can express the intention as well as identity of a vocalizer [6-10]. Intonation-sensitive brain areas have been reported in the human auditory cortex using subdural electrophysiology [11], and the ability to process and recognize pitch trajectory in vocalizations can be shaped by experience [12-14]. However, the temporal or spatial resolution of methods used in human studies are coarse compared to the rate at which neurons in the auditory system respond, as well as to the time scale of many intonations, leaving still unclear the neural mechanisms for tuning to the frequency modulations in natural sounds.

Frequency modulation encoding at the single neuron level in Core auditory cortex has been studied with directional sweeps or long duration sinusoidal modulations [15-20]. Offset responses are particularly selective for linear sweep direction [21], while entrained periodic firing can encode the onsets of the multiple cycles in up to 20 Hz sinusoidally modulated tones [15]. However, many natural tonal sounds have complex modulations over durations as short as a single cycle, and whether neurons show any tuning to the acoustics of these brief trajectories is not known. Furthermore, neither the ability of individual neurons in secondary auditory cortex (A2) to respond to such frequency modulations, nor the role that sound experience with behaviorally meaningful frequency modulations might play in shaping Core or secondary auditory cortical responses has been investigated in great depth.

Here we parameterized synthetic frequency modulations with both linear and sinusoidal components (sFM) to explore auditory cortical tuning to the acoustic parameters of a short sound’s frequency trajectory. We then applied this model to uncover experience-dependent sensitivities to frequency modulations in a natural vocalization paradigm, the mouse maternal model. Mice come to recognize pup ultrasonic vocalizations (or USVs) after maternal experience, and display preferential approach towards pup USVs over neutral sounds [22-24]. We conducted head-fixed, awake single unit (SU) electrophysiology in both maternal and non-maternal females, and presented sFM with systematically varied parameters to probe plasticity in frequency trajectory sensitivity across natural experience and auditory region. We found tuning to frequency modulation amplitudes less than 1/24^th^ octave, a much finer degree of FM sensitivity than previously uncovered. We then discovered an enhanced prevalence of Offset responses to natural USVs after maternal experience, specifically in A2. In maternal A2 SUs, a bias emerged in both On and Offset responses that favored vocalizations with pup-like sFM parameters. This bias coincided with a shift in tuning of the maternal A2 towards frequency trajectory parameters that were more characteristic of the pup USV category. Taken together, this work reveals how Offset tuning to frequency trajectory in A2 plays a key role in natural sound category learning.

## RESULTS

### Auditory cortical neurons can be tuned to subtle frequency modulations

Auditory neurons are often characterized with brief pure tones (PT) varying in frequency and intensity, and show tuning to a best frequency (BF) and best attenuation in this synthetic space (**Fig. 1A**). We were interested though in whether neural responses depend not just on the static spectrum of a tone, but also on small modulations in a sound’s frequency, as often found in natural sounds [25]. To this end, we tested single frequency sounds with the same 60 ms duration as used for pure tone tuning, but with sinusoidal frequency modulations (see Methods) parameterized by a spectral amplitude (A_fm_) and a temporal modulation frequency (f_fm_). We found that around a SU’s BF, frequency deviations smaller than the typical spectral width of pure tone tuning curves [26-28] often could drive better responses compared to the pure tone BF itself (**Fig. 1B**). The example SU had a peak in A_fm_ tuning at 1/10 octave (f_fm_ = 50 Hz), with an evoked spike rate more than twice that predicted from just integrating the pure tone excitatory tuning curve over the same spectral range (**Fig 1B**, green line). Even after doubling the duration of the sound (**Fig 1C**), so that spiking was seen not only after sound offset (Offset Resp), but also while the sound was still on (On Resp), this SU remained tuned in the A_fm_ parameter for a nonzero spectral amplitude. Intriguingly, larger A_fm_ values reduced firing rates from the peak, suggesting that this SU’s tuning would not be explainable simply by its excitatory sensitivity to the brief sound’s static spectrum.

**Figure 1.**
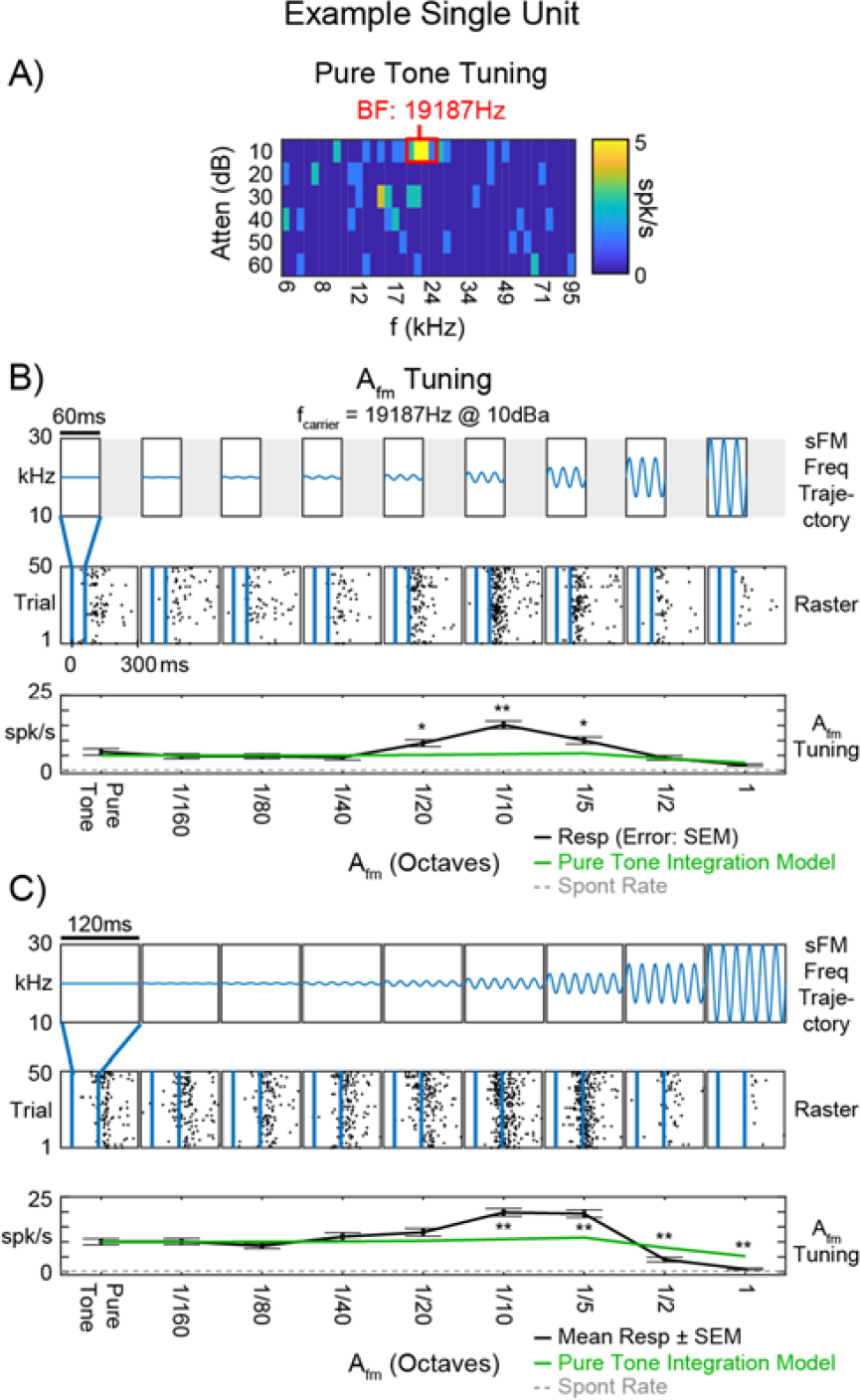
Example SU tuning to Afm in sFM sounds varied around BF. **A)** Pure tone tuning response area for SU 2981. The SU’s absolute spike rate is represented by the heat map color, with hotter colors representing higher spike rate. The SU’s best responding area is indicated by a red box, with a BF of 19187 Hz. **B)** A_fm_ tuning to 60 ms sFM varied around this SU’s BF. **Top**: Schematic frequency trajectories of each stimulus, with all other parameters fixed at: f_fm_ = 50 Hz, *φ* = 0, f_0_ = BF (19187Hz), f_slope_ = 0 Hz/s, dur = 60ms. A_fm_ is varied in logarithmic steps from 0 to 1 octave. **Middle**: Raster responses to stimuli delivered within the vertical blue lines. Individual spikes depicted by black dots. **Bottom**: Mean response tuning curve (black), with error bars representing standard error of the mean (SEM). Spontaneous rate (dotted gray line) and rate predicted from integrating the pure tone (PT) tuning curve (green) are also shown. *p<0.01; **p<0.0001 Bonferroni corrected t-test. **C)** A_fm_ tuning to 120 ms sFM varied around the SU’s BF. Same as in B, instead with the A_fm_ tuning stimulus using a duration of 120 ms.

Such results led us to ask whether SUs might instead be sensitive to how a sound’s frequency content unfolds in time, even for relatively short stimuli. We therefore played 60 ms sFM stimuli while varying not only A_fm_ around the pure tone BF, but also f_fm_, which governs how quickly the sound’s frequency traverses the spectral range across time. Looking separately at On and Offset Resp components for a different example SU (**Fig 2A**) while we played back these sFM stimuli (**Fig 2B**), Offset firing significantly increased as the tone’s frequency crossed the same A_fm_=1/13 octave range at slower f_fm_ rates, an effect that also held at neighboring A_fm_ values. However, tuning was different for On and Offset Resp components (**Fig 2C**), despite being similar for static pure tones (**Fig 2A**). Since the frequency trajectory immediately preceding the On and Offset Resp differ at the start and end of a modulated sFM sound, but are the same for a static tone, On vs. Offset tuning differences would be expected if the SU were sensitive to the local frequency trajectory. We therefore asked next whether such sensitivity can be found more generally across the auditory cortex by analyzing both On and Offset Resp to sFM stimuli for a larger population of SUs.

**Figure 2.**
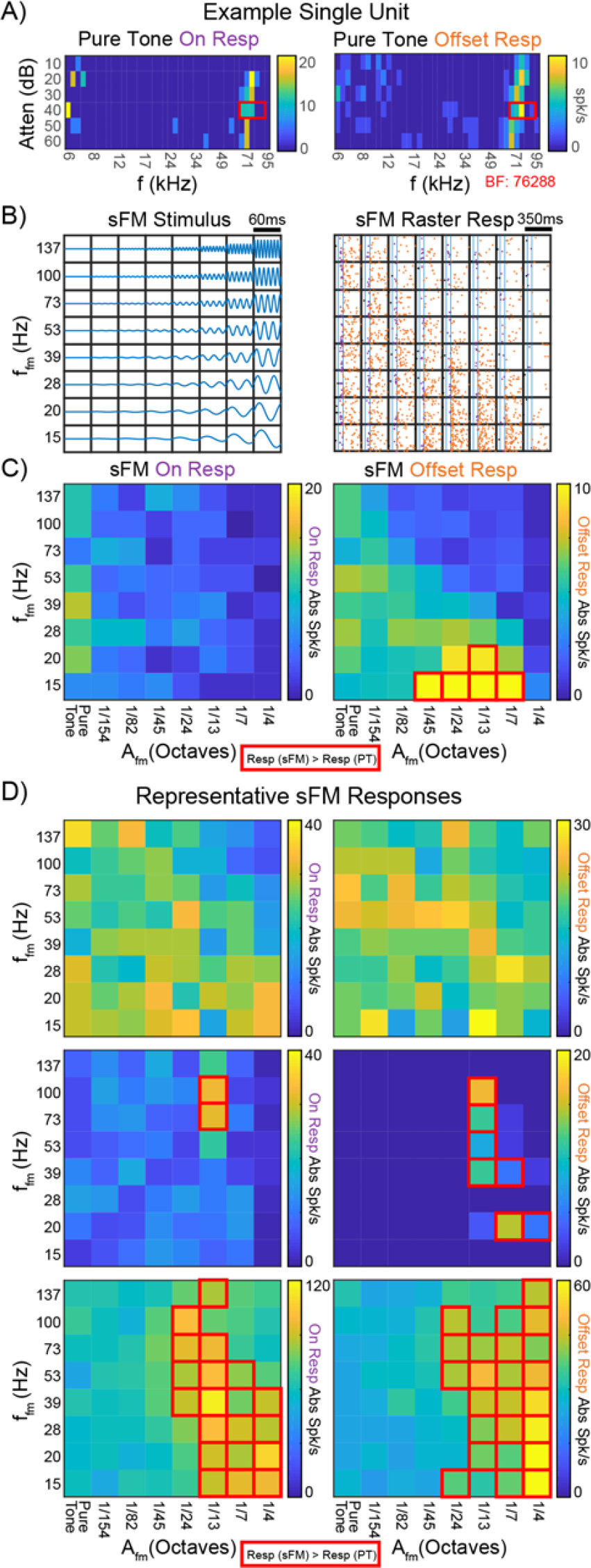
Example SU tuning to Afm x ffm in sFM sounds varied around BF. **A)** Pure tone tuning response areas for the On response (left) and Offset response (right) of SU 3009. The SU’s absolute spike rate is represented by the heat map color. The SU’s best responding areas for their On and Offset Resps are indicated by red boxes, with a BF of 76288 Hz. **B) Left**: Schematic frequency trajectories of the 60 ms duration A_fm_ x f_fm_ tuning stimulus. **Right**: Raster responses of the SU, with the purple dots representing individual spikes falling within the On Resp window, and the orange dots representing those in the Offset Resp window. **C)** Heat map representing the firing rate of the SU within the On Resp window (**Left**) and Offset Resp window (**Right**). In each grid, the left-most column, where A_fm_ is 0, consists of 8 sets of pure tone trials (25 trials each), and thus reflects the inherent variability in responses to the identical pure tone (PT) sound. Red boxes outline those stimuli that evoked a significantly higher spike rate compared to the PT spike rate (p<0.05, Bonferroni corrected Wilcoxon Rank Sum). **D)** On and Offset Resp heat maps for A_fm_ x f_fm_ stimuli in the case of three other example SUs (**Top**: SU 3164, **Middle**: SU 3064, **Bottom**: SU 3010). Red boxes same as in C.

### SU Offset Resps are more likely than On Resps to prefer modulated sFM frequency trajectories

We assessed the On and Offset Resp to sFM around BF across a population of n=61 Core (n=44) and secondary (A2, n=17) auditory cortical SUs, of which n=52 had an On Resp to at least one of these stimuli, and n=50 had an Offset Resp. SUs responded in varied ways in both their On and Offset Resp’s (**Fig 2D**). For some SUs, sFM sounds elicited spike rates (either On or Offset) that were comparable to pure tone responses (**Fig 2D**, top row) for all the stimuli within our sFM parameter space; these were considered non-tuned for sFM around BF. Many SUs, however, exhibited strong preferences either for sFM within narrow regions of combined, non-zero A_fm_ and f_fm_ (**Fig 2D**, middle row, red lines outline sFM stimuli that elicited a response significantly larger than for pure tones), or for larger ranges of A_fm_ and/or f_fm_ (**Fig 2D**, bottom row). Only 7/61 SUs (3/52 On, 4/50 Offset) showed significantly suppressed spiking for one of the sFM stimuli compared to pure tones, and thus were not considered in detail further. An excitatory preference for sFM could be seen in a SU’s On and/or Offset Resp. While larger frequency modulations typically elicited more responses that were significantly higher than pure tone responses, A_fm_ as low as 1/24^th^ of an octave (and on rare occasions even less) could modulate spiking in both the On and Offset Resp (**Fig 3A**). To determine which response type was more sensitive to sFM, we measured how many of the SUs with that response had at least one sFM stimulus that elicited a significantly greater spike rate than for the BF pure tone (p<0.05 Bonferroni corrected Wilcoxon Rank Sum). We found that Offset Resps were significantly more likely than On Resps to have a preferred sFM different from pure tone (40% vs. 17%, p<0.05 Fisher’s Exact, **Fig 3B**), as might be expected if a SU were responding to a sound’s frequency trajectory and not just its spectrum. In fact, even for those SUs with a sFM-preferring On Resp, those On Resps were typically more sustained, tonic, or late-onset. Hence, more of a sound’s history than just its onset can be integrated to affect the neural response, as would be the case for Offset Resps. An additional control further confirmed a neural sensitivity to short frequency trajectories as they unfold over time, and not just a stimulus’ spectrum. We played back to some SUs a narrowband noise that was bandwidth-matched to each of the sFM stimuli in our 8×8 A_fm_ x f_fm_ grid. SU Offset Resp’s were significantly higher for sFM compared to their matched noise (**Fig 3C**, left, p<0.01 Paired Wilcoxon Signed Rank). Taken together, these data suggest that Offset Resps are particularly sensitive to frequency trajectory variations present in short sFM stimuli.

**Figure 3.**
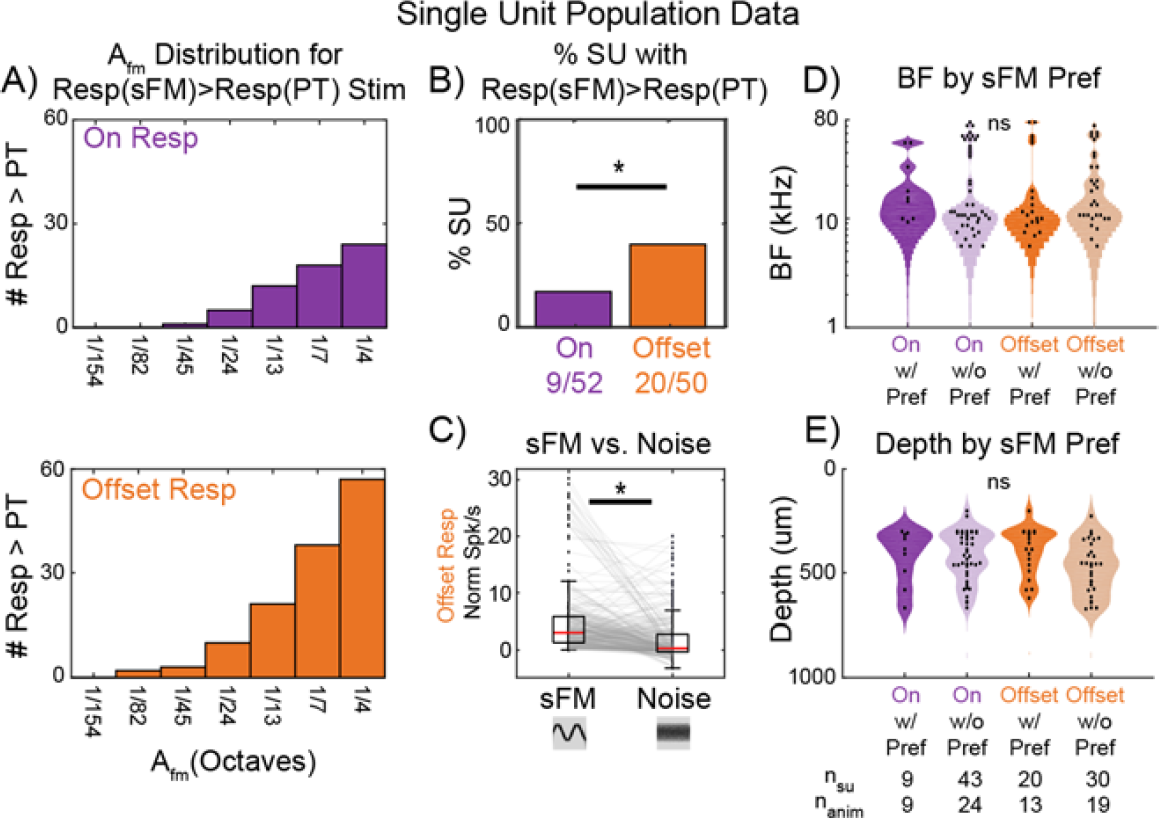
sFM tuning across the neural population. **A)** Histogram of responses significantly modulated by subtle sFM. Each panel shows the distribution of A_fm_ values across SUs and A_fm_ x f_fm_ combinations for sFM stimuli that elicited a significantly greater response than the SU’s pure tone response for both On Resps (**Top**) and Offset Resps (**Bottom**). Significance was based on each SU’s statistical testing (see red boxes in **Fig 2C**). **B)** Offset Resps are more likely to be significantly modulated by sFM trajectories. The proportion of SUs with an On Resp that prefer sFM stimuli over pure tone was significantly lower than the proportion of SUs with an Offset Resp that prefer sFM stimuli over pure tone (p<0.05, Fisher’s Exact Test). **C)** The sFM trajectory elicits stronger Offset Resps than noise bursts with matched bandwidth (left, n_stim*su_ = 280, n_su_ = 40, n_anim_ = 21). * p<0.05, Paired Wilcoxon Signed Rank. **D)** SU BF distributions did not differ depending on whether the SU showed a (non-PT) sFM preference or not in their On or Offset Resp (ns, Kruskal Wallis). **E)** SU depth distribution did not differ depending on whether the SU showed a (non-PT) sFM preference or not in their On or Offset Resp (ns, Kruskal Wallis).

### sFM-sensitive neurons are found across the hearing range and across cortical depths

SUs that showed a preference for sFM over pure tones had BFs varying across the entire mouse hearing range from 6-80kHz, with BF distributions that were not significantly different between On or Offset Resps (**Fig 3D**). These BF distributions were also comparable to that seen for SUs without a preference for sFM. Functionally, this implies that frequency trajectory sensitivity could be a neural mechanism to attune to a wide variety of behaviorally relevant sounds across any part of the mouse’s hearing range. Furthermore, the distribution of cortical depths between SUs with or without sFM preference also did not differ (**Fig 3E**); SUs showing preference for sFM spanned the entire range of cortical depths that were sampled (200-700μm).

Together, these results suggest a general ability of auditory cortical neurons to attune to frequency trajectory parameters in any sound, especially in their Offset Resps. Presumably such sensitivity to subtle frequency modulation, which is prevalent in many natural sounds, would be particularly useful in accurately representing features of acoustic categories that must be behaviorally discriminated, even when those sounds overlap in their overall spectrum. If so, we would expect that experience to make a sound category meaningful could drive neural plasticity in the tuning. To address that possibility, we focused next on characterizing trajectory sensitivity to ultrasonic vocalizations (USVs), which are ethologically relevant to mice, and investigated whether and where neural plasticity in USV trajectory sensitivity emerges after the behavioral relevance of a specific USV category is acquired through experience.

### SUs show heterogeneity in timing of responses to USVs

Adult female mice come to recognize USVs emitted by isolated pups after maternal experience, as demonstrated by their preference to approach a pup USV over a neutral sound [23, 24]. Adult males emit another category of USVs that is also meaningful for females, and these overlap the pup USV frequency range. Several studies have demonstrated that learning the behavioral relevance of pup USVs leads to both inhibitory and excitatory plasticity in the response to USVs within Core auditory cortex [29-34], and that the ability of a specific subset of Core neurons to discriminate between the pup and adult USV category is heightened after experience [32]. Importantly, this neural discrimination is apparent even for pup and adult USVs matched in their onset frequencies and durations. This raises the critical question of how neurons are able to differentiate these brief sounds, and whether they might be attuning to the frequency trajectories of specific call types to enable discrimination.

Given the importance of Offset Resps for attuning to frequency trajectories, we investigated this question by first determining the degree to which SUs in both Core and secondary (A2) auditory cortex exhibit Offset Resps to USVs. We used a stimulus set of n=36 curated, natural pup and adult USVs with matched onset frequency and duration properties (**Fig 4A**). Following previous work [30, 32], we classified SUs from maternal and nonmaternal animals (see Methods) as call-excited, call-inhibited, or call-nonresponsive (NR). Notably, a sizable fraction of neurons we recorded were not responsive to any USVs (NR n=449/720), evidencing our attempts to reduce unintended recording bias against those SUs that have lower spontaneous firing rates or highly selective responses. SUs that were purely call-inhibited, which have been studied elsewhere [24, 30], were not considered further here.

**Figure 4.**
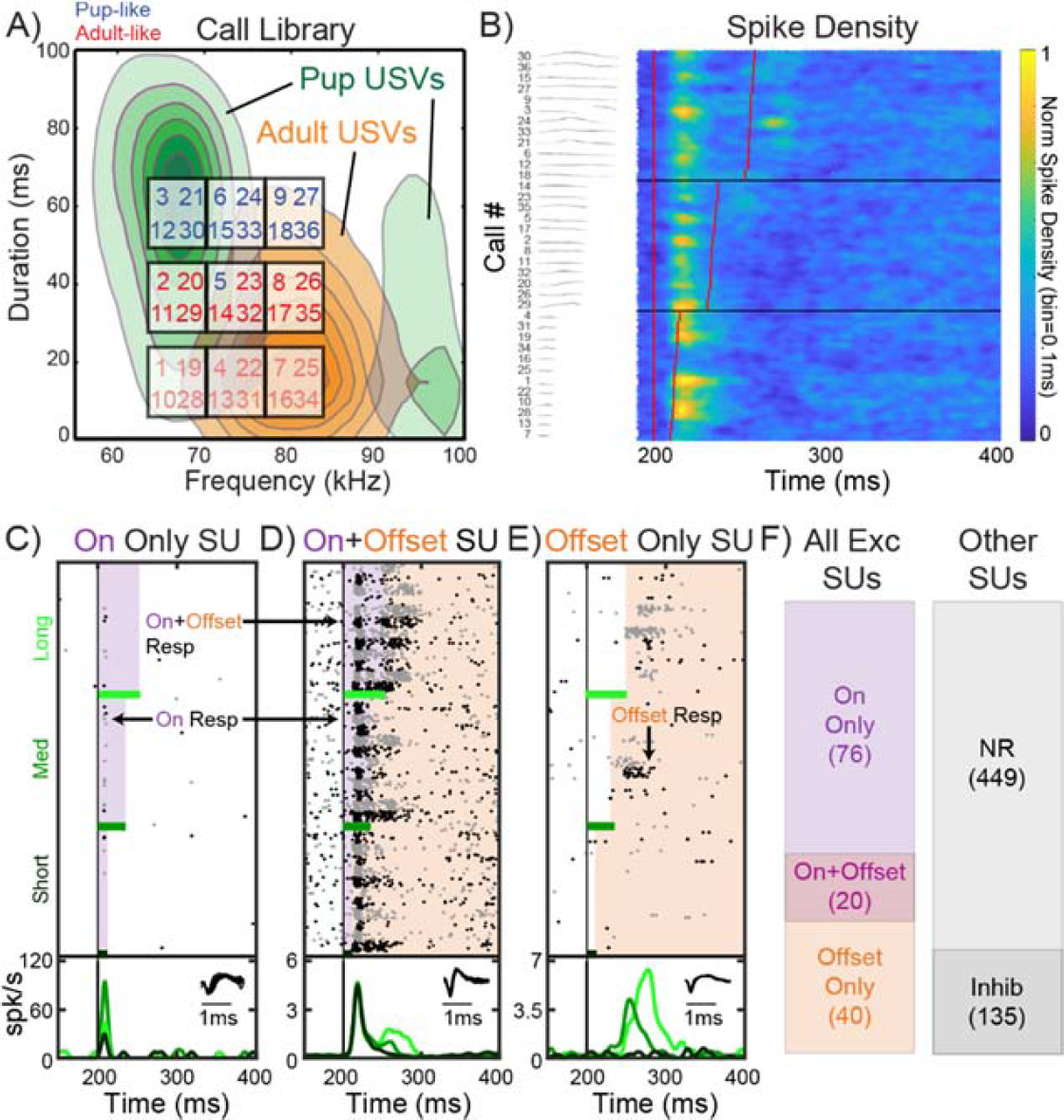
Classification of On and Offset Resps to USVs. **A)** Onset frequency and duration of the ground-truth pup (1-18) and adult (19-36) USVs in our playback library. The underlying distributions for pup (green) and adult (orange) USVs are shown as contours of increasing likelihood for calls of each ground-truth category to have specific duration and frequency values. Because our calls were chosen to systematically vary in duration and frequency, some actual pup calls are acoustically more similar to adult calls, and vice-versa. This is apparent from the logistic regression model that allowed us to relabel the USVs as pup-like (blue) or adult-like (red) depending on whether their frequency trajectory parameters were more predictive of being a pup or adult call, respectively. Note that the bottom row of (adult-like) USVs are shown muted because these are withheld in some analyses that required evoked responses to be definitively classified as On or Offset, and those calls are too short to do so accurately. **B)** Spike density plot depicting overall population responses to USVs across all call-excited SUs (n=136, n=55 animals), sorted by increasing USV length. The duration of each USV is indicated by the vertical red lines, starting with playback at 200 ms. Left column shows the USV spectrograms. **C)** Representative SU with only On Resps to several USVs. Rasters alternate between black and gray to delineate the trials of adjacent calls. Purple shaded area represents the On Resp window. Bottom panel shows the Peristimulus Time Histogram (PSTH), which pools responses across all calls within the same duration group: short, medium and long (rows in **Fig 4A**). Inset shows the SU’s spike waveform. **D)** Similar to **C**, but for a SU with both On Resps and On+Offset Resps to different USVs. Orange shaded area represents the Offset Resp window. **E)** Similar to **C**, but for a SU with only Offset Resps to several USVs. **F)** Overall classification of recorded SUs by their USV-evoked response characteristics. Left, breakdown of call-excited SUs. Right, breakdown by non-responsive (NR) and call-inhibited (Inhib), which were not included in subsequent analyses looking at excitatory tuning to USV features.

Call-excited responses (n=136) across Core and A2 were pooled to create a spike density plot for USVs, revealing evidence in the population data for Offset Resps to individual calls (**Fig 4B**), especially for the longest USV durations. We therefore classified each call-excited SU into one of three sub-categories. On Only SUs had an On Resp to at least one of 36 calls, but no Offset Resps to any (**Fig 4C**). On+Offset SUs had an On Resp to at least one of 36 calls, and an Offset Resp to at least one call as well (**Fig 4D**); this could include On+Offset Resps for the same call. Finally, Offset Only SUs had an Offset Resp to at least one of 36 calls, but no On Resps to any (**Fig 4E**). Most SUs we recorded were On Only (**Fig 4F**), but about 44% of call-excited SUs had some form of Offset Resp, falling into either the On+Offset or Offset Only groups. Hence, although there was considerable heterogeneity in when auditory cortical SUs increased their firing rate in response to USVs, a sizeable portion did so after the end of at least one of the calls, opening the possibility that these excitatory Offset Resps were sensitive to the preceding frequency trajectory in calls.

### Strength of On Resps to USVs decreases with experience

We next reasoned that if this were indeed the case, then as particular categories of calls become more behaviorally relevant, neural responses would change to improve the coding of newly meaningful frequency trajectories. In numerous paradigms, behaviorally meaningful experience learning about sounds induces different forms of auditory cortical plasticity [29, 35, 36]. To determine whether maternal experience caring for pups might alter tuning for natural USV frequency trajectories, we analyzed whether this experience affects the strength of USV-evoked On and Offset Resps across auditory cortical fields (Core: A1, AAF, Ultrasound Field UF, versus A2) by comparing maternal animals to nonmaternal animals. We calculated On and Offset Resp spike rates on a per-call basis and normalized by subtracting a SU’s spontaneous rate. We found that *on average* in the Core, experience did not affect either On or Offset Resp spike rates (**Fig 5A**, top), although there was considerable variability. Interestingly though, in A2, maternal animals showed a significantly decreased On Resp firing rate (p<0.0001, Bonferroni corrected Wilcoxon Rank Sum), but we saw no changes in evoked Offset spike rates (**Fig 5A**, bottom). When analyses were performed on a per-SU basis (see Methods) rather than a per-call basis, A2 still showed significantly decreased On Resp spike rates in maternal animals (p<0.01, Wilcoxon Rank Sum).

**Figure 5.**
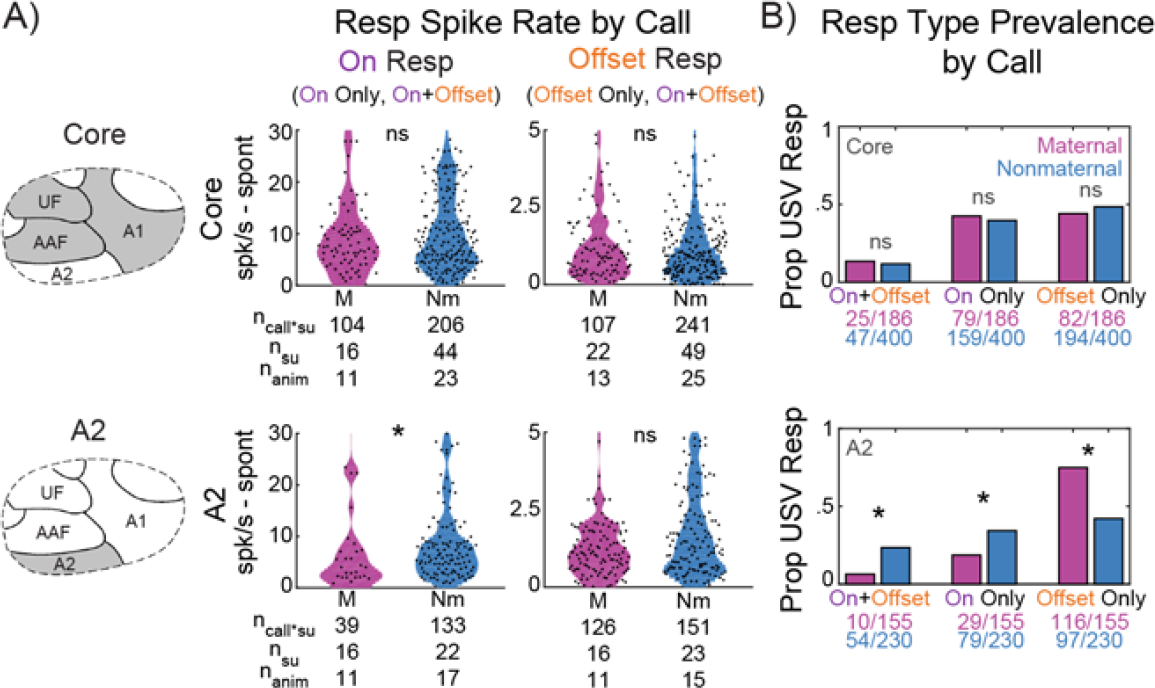
On and Offset Resps to USVs differs across field and maternal experience. **A)** Response spike rates on a per-call basis divided by auditory cortical field – Core (**left**, **top row**) and A2 (**left**, **bottom row**) – and by On (**middle**) and Offset (**right**) Resp. Spike rates are normalized by subtracting an SU’s spontaneous rate. SUs from maternal animals shown in magenta and from nonmaternal animals shown in cyan. * p<0.0001, Bonferroni corrected Wilcoxon Rank Sum. **B)** Prevalence of each type of Resp on a per-call basis, divided by auditory field – Core (**top**) and A2 (**bottom**). * p<0.0001, Fisher’s Exact Test.

### Prevalence of Offset Resps to pup USVs increases with experience

Aside from the strength of the evoked response itself, changes may also arise in the proportion of On or Offset Resps being elicited by USVs. To address this, we computed, on a per-call basis, the proportion of all call-excited responses from maternal or nonmaternal SUs that showed an On Only, On+Offset, or Offset Only Resp to the call. In the Core, we found no change in these proportions after maternal experience (**Fig 5B**, top left). However, in A2, the prevalence of Offset Only Resps increased significantly (p<0.0001, Bonferroni corrected Fisher’s Exact), while On Only and On+Offset Resps decreased significantly (**Fig 5B**, bottom left, p<0.0001 Fisher’s Exact). Even on a per-SU basis using the SU classification from **Fig 4C-E**, Offset Only A2 SUs were still significantly more prevalent in maternal animals (p<0.005, Bonferroni corrected Fisher’s Exact).

Hence, maternal experience altered both On and Offset Resps at a population level in A2 more so than in Core auditory cortex. Even though we found Offset Resps in both Core and A2, A2 was particularly plastic in terms of how often calls evoked Offsets, albeit not in how strongly they evoked spiking if they did respond. The observed plasticity thus tilted the balance in A2 away from responding at the beginning of these short sounds towards signaling as a population more at the calls’ terminations, which could better enable frequency trajectory sensitivity. If this were indeed the case though, we expect that Offset responses in A2 would become better attuned to the frequency trajectories of natural USVs. To determine this, we first characterized the acoustic parameters of *full* frequency trajectories in natural USVs, going beyond our previous characterization of just onset and duration properties [32, 37].

### Combined sinusoidal with linear frequency modulation sufficiently models frequency trajectories in natural USVs

We fit a large library [37] of 57,929 pup USVs and 10,353 adult USVs to a six parameter sFM model that added linear to sinusoidal modulation to accurately describe the frequency trajectories of USVs (**Fig 6A**, see Methods). The mean square error for the fit to each call was relatively small, averaging 639 Hz, compared to the 60-80 kHz range of the USVs themselves (**Fig 6B**, left). Both Offset Resps and full responses (data not shown) elicited by these sFM model USVs were highly correlated with responses evoked by their paired, natural USVs with the same amplitude envelope (**Fig 6C**, Spearman rho=0.92, p<0.0001). Hence, our parameterized frequency modulation largely captured the frequency trajectory features in USVs that drive SU firing.

**Figure 6.**
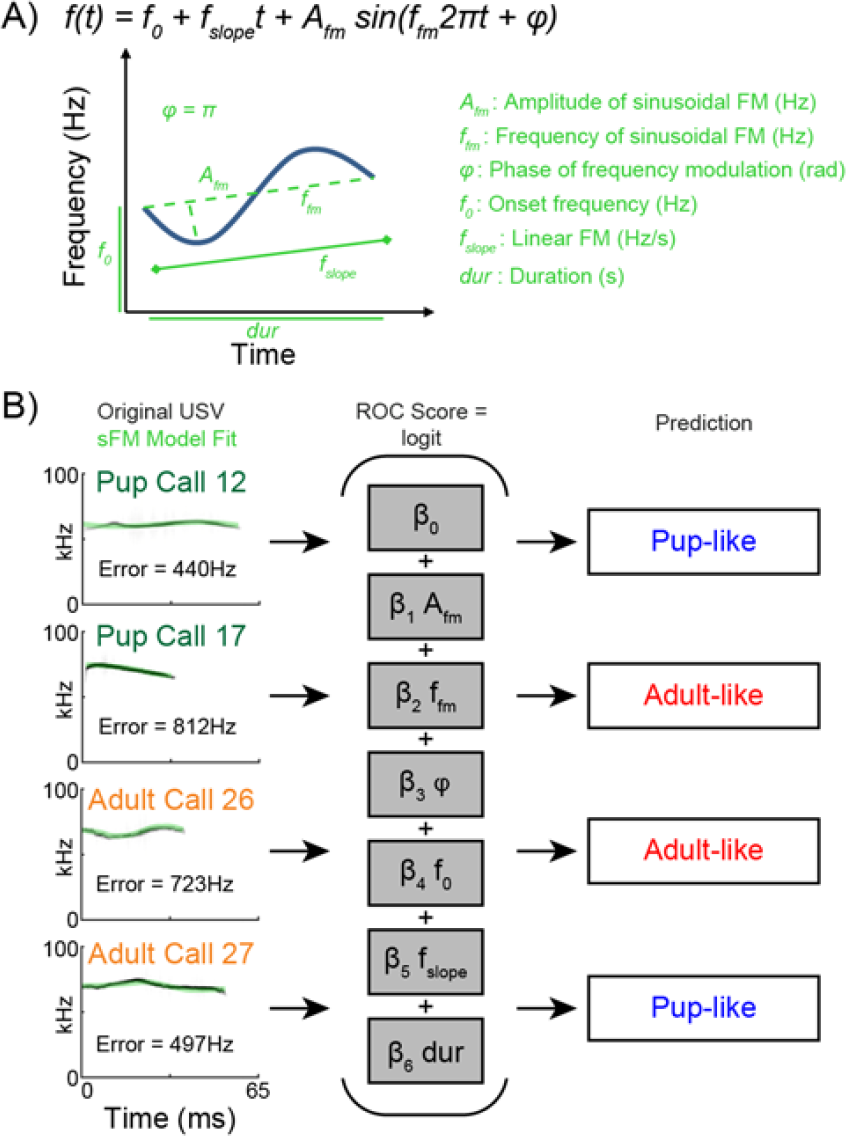
sFM models of USV trajectories can discriminate call categories. **A)** Schematic illustrating the sFM model equation. **B)** Nominal logistic regression modeling to classify USVs. (**Left**) Frequency trajectories of sFM models (light green) of ground-truth pup and adult USVs, overlaid on the spectrograms of the original USVs (grayscale). For each call, a set of six parameters was generated, which was then used to predict the call’s likely category (pup-like, blue; adult-like, red) using a nominal logistic regression model. Error was defined as the Sqrt(sum squared error / number of samples). Model parameters: β_Afm_ = −1.6e-5; β_Ffm_ = −2.4e-3; β_φ_ = 8.0e-2; β_f0_ = - 9.4e-6; β_fslope_ = −2.1e-8; β_dur_ = 3.4e-2; Model χ^2^ = 7.2e3, p<0.0001. **C)** Comparison of Offset Resp spike rates evoked by an original, natural USV versus its sFM model. There is a high correlation between responses (Spearman rho=0.92, p<0.0001). **D)** Control stimulus set comparing Offset Resps to sFM models of USVs versus just the linear FM component of the model, and the spectrally matched noise (right, for sFM vs lFM: n_stim*su_ = 35, n_su_ = 20, n_anim_ = 13; for sFM vs Noise: n_stim*su_ = 19, n_su_ = 14, n_anim_ = 8). * p<0.01, Paired Wilcoxon Signed Rank. **E)** Performance of the nominal logistic regression model according to an ROC analysis in correctly classifying ground-truth pup calls as pup-like (true positive) and in incorrectly classifying ground-truth adult calls as pup-like (false positive). Threshold (gold) represents the cutoff at which the true positive rate is maximized and the false positive rate is minimized. The model performs above chance (Area Under Curve = 0.76, p<0.0001 via Boostrap analysis, N=1000).

To address whether SUs may be attuned to the natural USV trajectories, we considered two control stimuli. In a subset of SUs, we presented not only our full sFM model of a USV in our library, but also a narrow-band noise model whose power spectrum was exactly matched to the calls’ (see Methods), and/or just the linear frequency modulation (lFM) component of our sFM model. The latter control addressed the possibility that responses to spectrally matched noise were simply suppressed by the noise spectrum engaging a SU’s inhibitory sidebands [38]. In all cases, we applied the amplitude envelope of the paired natural call. We found that Offset Resp’s for our full sFM+lFM model were preferred over both the spectrally matched noise and the Lfm component alone (**Fig 6D**, right, p<0.01 Paired Wilcoxon Signed Rank). Hence, SUs were indeed sensitive to how our short USVs changed over time from one frequency to another, consistent with what we observed more generally for sounds across the mouse hearing range (**Fig 3B**). We next investigated whether this sensitivity could potentially allow SUs to attune to those frequency trajectory parameters that differentiate one meaningful acoustic category from another.

### Specific sFM parameters distinguish stereotypical pup-like and adult-like USVs

To determine whether sFM frequency trajectory parameters are distributed so as to acoustically distinguish individual pup USVs from adult USVs, we first applied a nominal logistic regression model to our calls to find parameter ranges for the most stereotyped calls in each category (see Methods). For each of the six sFM parameters, we determined their optimal weight for calculating all calls’ Scores in order to maximize accuracy in predicting ground-truth “pup” and “adult” USV labels (**Fig 6B**, right). Due to multi-dimensional overlap of pup and adult USVs in acoustic space, as typically seen for natural vocal categories [37], our call classification model was not perfect. Nevertheless, it performed significantly above chance according to a Receiver Operating Characteristics (ROC) analysis (**Fig 6E**, p<0.0001, Area Under Curve AUC=0.758, 95% Confidence Interval (CI): 0.754 – 0.762, Bootstrap N=1000). The best performing Score threshold was 0.869, where a score >0.869 indicated the call would be classified as having a “pup-like” set of parameters, and all others as “adult-like.” For this choice, the model’s sensitivity (true positive rate) was 62.0%, and 1-specificity (false positive rate) was 16.8%.

Using this model, the overall distributions of our sFM parameters for “pup-like” and “adult-like” calls (**Fig 7A**) revealed that pup-like calls had systematically lower f_0_, higher A_fm_, longer durations, and higher φ than adult-like calls, suggesting that these parameters were particularly helpful for differentiating stereotyped pup from adult calls. Among the 18 ground-truth pup USVs in our curated set of played back calls (**Fig 4A**), 7 were classified as pup-like by the ideal observer (calls 3, 5, 6, 9, 12, 15, 18 in **Fig 4A**), indicating that these were good exemplars of what are the most stereotypical pup calls. Notably, all 6 short duration, ground-truth pup calls (calls 1, 4, 7, 10, 13, 16) were classified as adult-like, presumably because their short duration was more stereotypical of adult calls. We originally chose our curated set to uniformly span acoustic space (in duration and onset frequency), so it is not surprising that some ground-truth pup calls would have sFM parameters falling into the adult-like space, or vice-versa. Nevertheless, our acoustic analysis allowed us to identify those of our 36 exemplars that were most stereotypically pup-like or adult-like in their trajectory, so that we could examine how auditory cortical SUs respond to these features.

**Figure 7.**
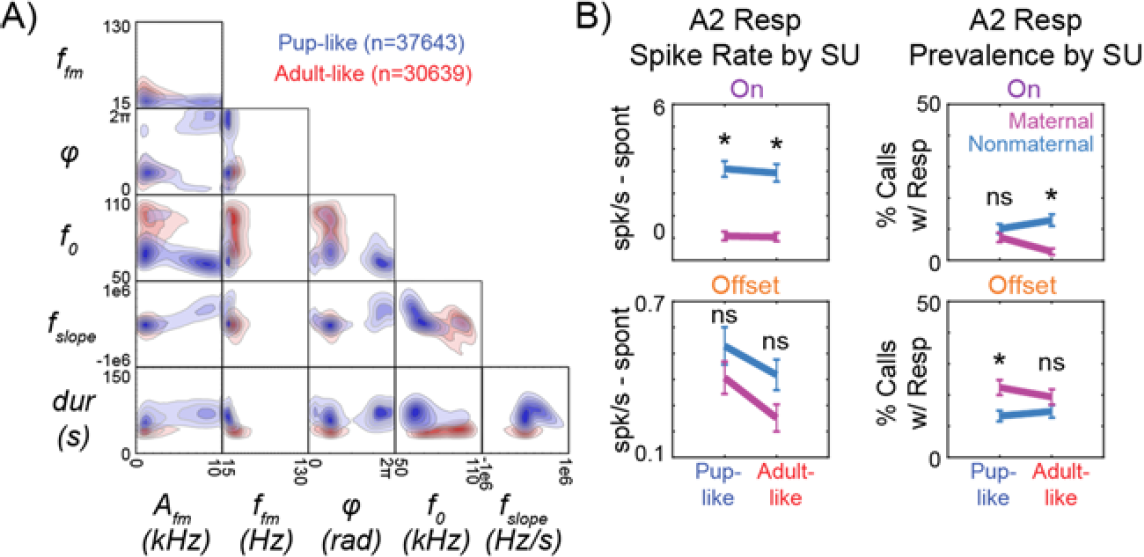
Maternal plasticity in A2 responses to USVs depends on whether USV trajectories are pup-like or adult-like. **A)** Distribution of the six sFM parameters for pup-like (blue) and adult-like (red) USVs. Separation between categories is apparent for A_fm_, duration, phase (*φ*), and onset frequency (f_0_). **B)** SU spike rate responses (**left**) and prevalence (**right**) to our curated (medium and long duration) USVs during the On Resp window (**top**) and Offset Resp window (**bottom**), separated by USVs that are pup-like versus adult-like, as classified by the logistic regression model. * p<0.05 Bonferroni corrected Fisher’s Exact Test.

### A2 On and Offset Resp plasticity depends on stereotypical features of pup-like and adult-like USVs

With the finer acoustic classification, we next reconsidered the decreased On Resp spiking and increased Offset Resp prevalence for USVs in A2 of maternal animals (**Fig 5**). We found that while suppressed On Resp spiking in maternal animals occurred universally for both pup-like and adult-like USVs (p<0.05, Bonferroni corrected Fisher’s Exact; **Fig 7B**, left), the prevalence of On Resps *decreased* specifically for adult-like calls, while the prevalence of Offset Resps *increased* specifically for pup-like calls (p<0.05; **Fig 7B**, right). Hence, in the maternal A2, SUs become more likely to show Offset Resps precisely for calls that have frequency trajectories that are more stereotypically pup-like, highlighting a new form of plasticity in Offset Resps that depends on the statistical properties of a newly meaningful sound category.

### Resp tuning for A_fm_ shifts in the maternal A2 to enhance stereotypical pup USVs

Finally, we asked whether A2 SUs’ responses change in their ability to encode frequency trajectory parameters after maternal experience via changes in their parameter response tuning curves. In a subset of SUs, we selected the curated natural USV that elicited the best response, and then systematically varied A_fm_ around the fixed values of the other 5 sFM parameters that modeled this best call (**Fig 8A**). The resulting tuning curve for A_fm_ (**Fig 8B**) was fit with a Gaussian to identify both the best A_fm_ and the point of greatest slope (Max slope).

**Figure 8.**
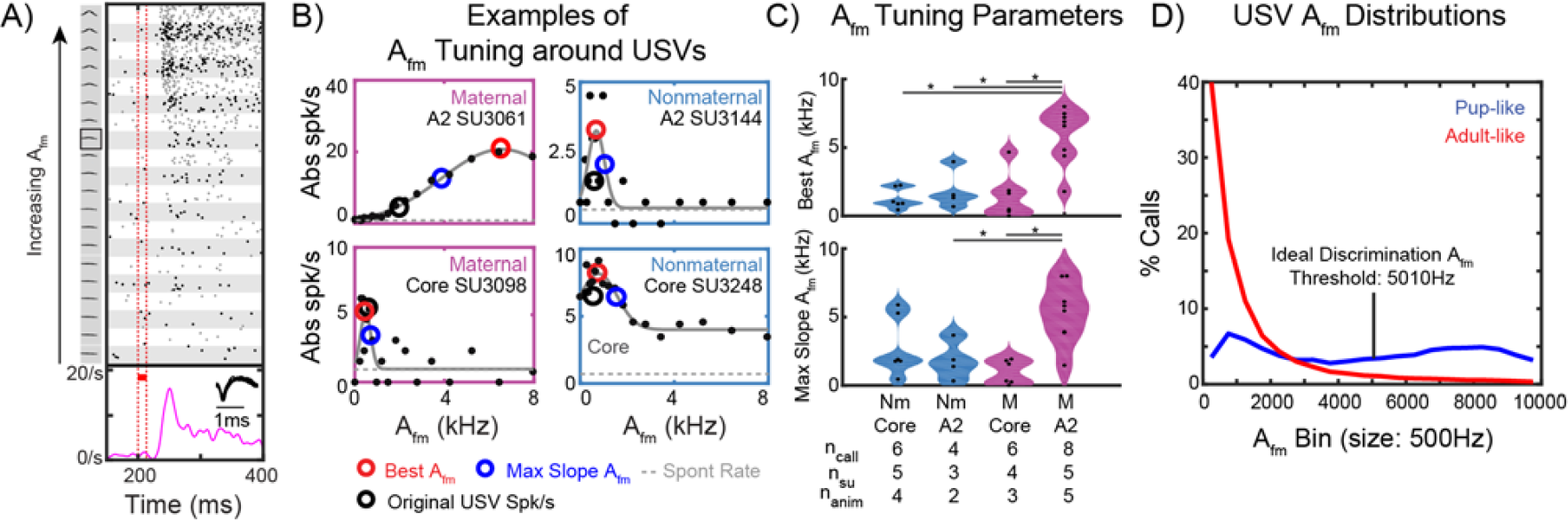
Afm tuning around USVs changes in A2 with experience. **A)** Raster responses to sFM models with A_fm_ varying around an example SU’s best call (boxed spectrogram in left column). Tuning stimuli, presented during the period demarked by the red lines, include 19 different sFM models with A_fm_ varying from 0 to 8000 Hz, which spans the range of natural USV A_fm_ values. The other sFM parameters are fixed at values matching the best USV. PSTH shows pooled spike rate responses to all stimuli, highlighting the responses after the Offset of the sounds. **B)** Example A_fm_ tuning curves of four different SUs, based on the full responses. Two SUs from A2 (**top**) and two from Core (**bottom**) are shown from both maternal (**left**) and nonmaternal (**right**) mice. Responses are fit to a Gaussian, whose peak is denoted by a red circle, and point of maximum slope by a blue circle. Spike rate in response to the original USV is denoted by a black circle. Spontaneous rates indicated by a dotted gray line. **C)** Best A_fm_ (**top**) and Max Slope A_fm_ (**bottom**) values according to animal experience group (M = maternal, Nm = nonmaternal) and auditory field. *p<0.05 Tukey Kramer’s HSD. **D)** Histogram of A_fm_ in either the pup-like (blue) or adult-like (red) USVs, as a proportion of the calls in each respective category. The ideal observer discrimination threshold for pup-like versus adult-like calls, as determined by ROC, is indicated by the black line at A_fm_ = 5010Hz.

We found that responses were not always strongest for the specific natural call we played, but that SUs were nevertheless tuned in A_fm_. SUs in the maternal (M) A2 group generally preferred significantly higher A_fm_ values than those in nonmaternal (Nm) animals or in the Core (**Fig 8C**, top, on a per-call basis, p<0.005 Tukey-Kramer Honestly Significant Difference (HSD)). Moreover, maternal A2 SUs also had a significantly larger Max Slope A_fm_ compared to the maternal Core or nonmaternal A2 groups (**Fig 8C**, bottom p<0.05 Tukey-Kramer HSD). Despite stimuli for different SUs having different starting sFM parameter values since they were centered around different best calls, we did not find systematic differences in those other sFM parameter values between the SUs in each group. Furthermore, since in several cases an SU’s best call was one of those with short durations (**Fig 4A**, bottom row), we used the full response rather than just the Offset Resp, since we could not completely differentiate Offset Resps from delayed On Resps in these cases. However, within the entire cohort of n=24 SUs, very few responded during stimulus playback itself (n=2 maternal Core SUs, n=2 nonmaternal Core SUs and n=0 A2 SUs), and our results still held if we only considered responses after the stimulus ended. Our results also held on a per-SU and per-animal basis (using the average best A_fm_ value in each case). Importantly, if we instead compared the best A_fm_ for stimuli varied around a SU’s BF instead of around a USV, then we saw no differences across animal groups or region (p=0.22 maternal vs nonmaternal, p=0.59 Core vs A2, Wilcoxon Rank Sum, data not shown). Hence, our data demonstrate an upward shift in A_fm_ tuning of responses that is specific to tuning around USVs in maternal A2 SUs.

As a last step, we sought to gain some insight into why such tuning changes in A2 might be functionally useful for maternal animals in their encoding of A_fm_ in pup-like sounds. The A_fm_ value of a given USV was correlated with its pup-likelihood, as measured by the logistic regression Score, so that higher A_fm_ predicted higher pup-likelihood Scores (Spearman rho = 0.6766; p<0.0001). The distribution of A_fm_ across pup-like calls showed a larger proportion of high A_fm_ values compared to adult-like calls (**Fig 8D**). Using ROC analysis, we found that an A_fm_ value of 5009.7 Hz best discriminates pup from adult USVs (AUC=0.61574, p<0.0001; True Positive = 38.8%; False Positive = 13.15%). Interestingly, the mean Max Slope A_fm_ value of the maternal A2 group was 5329Hz, close to the ideal pup-like versus adult-like USV discrimination point (**Fig 8C**, bottom), whereas that for non-maternal A2 was only 1819Hz. Hence, our results suggest that with experience, responses in A2 shift their tuning in a key frequency trajectory parameter (A_fm_) in a way that could improve discrimination of a newly meaningful sound category from other natural sounds.

## DISCUSSION

Intonation contributes to the expression of emotion [4, 39], the conveyance of prosody in speech [40] or vibrato in music [41], and even the learning of speech sounds [42]. However, we still have a poor understanding of how the brain attunes to such complex, behaviorally relevant frequency contours. Here we showed that the strength of offset firing by auditory cortical neurons reflects its sensitivity to the precise frequency trajectory that a sound took before ending. Frequency trajectory fluctuations is as low as 1/24^th^ octave, much finer than previously realized could significantly modulate responses. Furthermore, offset responses exist in both primary and secondary auditory cortical fields. Importantly, those in mouse A2 are particularly plastic after experience with a meaningful sound category whose short pitch trajectories differentiate one vocal category from another. A specific increase in the prevalence of A2 Offset Resps to pup-like USVs in maternal mice who have cared for pups is accompanied by a retuning in the A2 sensitivity to ultrasonic frequency trajectories. Taken together, these results demonstrate that a secondary auditory cortical field’s offset firing provides a neural substrate for experience-dependent encoding of behaviorally relevant intonations, such as those in natural, emotional, communicative sounds [43, 44].

Offset responses in the central auditory system are of growing interest. Besides being generally implicated for detecting silent gaps in sounds [45, 46], we now know that compared to onset firing, sound offset firing is mediated through separate synapses in the cortex [47], tuned to different pure tone frequencies [21, 45, 47-49], and spatially localized in distinct ways [50, 51]. However, in part because earlier studies only used noise or constant tones, the tuning of offset responses to subtle modulations in a sound’s frequency trajectory had been overlooked. Here we found that if a neuron is excited by a particular frequency trajectory, that response cannot be explained just by a sound with the same static spectrum, or by a linear ramp from the initial to final frequency (**Fig 3C**, **6D**). In fact, many neurons’ responses are affected by changing the modulation rate without altering the spectral extent of modulation (**Fig 2**), highlighting the importance of the precise frequency trajectory in determining the response. The temporal integration of excitatory and inhibitory synaptic inputs elicited by a sound’s frequency trajectory is presumably critical for generating this tuning. Computational modeling [21, 45], beyond the scope of the current work, will likely be essential for clarifying how this tuning arises only after the sound ends, especially given the generally broad pure tone excitatory tuning of many mouse auditory cortical neurons [27, 28].

The encoding of frequency modulations by auditory neurons is of much general interest and has been studied previously in a number of ways. Sweep direction selectivity and sweep velocity preference were observed in primary auditory cortex using stimuli with linear or logarithmic unidirectional frequency modulation (e.g. [17, 18, 52, 53]). Phase-locked firing during sinusoidal frequency modulations in tones lasting on the scale of seconds has also been reported [15, 16, 20]. However, natural sounds often feature much more complex modulation than just unidirectional or sinusoidal sweeps alone, yet few studies have systematically explored such combined modulations. Our study was motivated to do so based on the acoustic analysis of mouse USV trajectories (**Fig 7A**), and the fact that while unidirectional frequency sweep responsiveness correlates with how well neurons respond to complex trajectories in vocalizations [54], such responsiveness measures do not capture whether neural activity is actually tuned locally to complex modulations (**Fig 1-2**, **8**). By approximating natural calls with our 6-parameter sFM frequency trajectory model (**Fig 6**) to explore tuning, we revealed not only narrow tuning around a neuron’s BF (**Fig 3**), but also graded neural firing as acoustic parameters were varied around natural USVs (**Fig 8B**). Narrow tuning in frequency modulation parameter space might therefore serve as an additional mechanism beyond so-called combination-sensitivity [55-57] to create sparse selectivity in some neurons for specific sound frequency trajectories (e.g. **Fig 4E**).

While most work on auditory cortical neural coding has focused on Core fields, much less is understood about the role of non-Core fields, and no studies to our knowledge have explored the systematic encoding of complex frequency modulation in A2. We observed tuning to sFM acoustic parameters in both Core and A2 neurons, with similar general characteristics, allowing us to combine those neural populations in presenting tuning around BF (**Fig 3**). This agrees with a recent report [51] using wide-scale calcium imaging to observe a strong Core as well as A2 Offset response to long-duration tones, particularly in the ultrasound range. In our study, in response to natural USVs, evoked On and Offset firing rates and prevalence among well-isolated, call-excited single units, were comparable between Core and A2 (**Fig 5**). Moreover, the ∼44% call-excited neurons with Offset responses (**Fig 4F**) we observed is similar to the proportion found in mouse medial geniculate nucleus (MGN) [45], especially in both the ventral and dorsal divisions, which project respectively to Core and A2 in the mouse [58]. Hence, extending what has been known for pure tone coding in subcortical auditory areas [19, 59], our results confirm that Offset responses are prevalent in both primary *and* higher order auditory cortical fields. Importantly, our study newly suggests that such responses throughout the auditory system may be particularly sensitive to frequency modulation.

In fact, by combining the mouse maternal model for naturally increasing the behavioral relevance of vocal categories [60] with a parameterization of USV frequency trajectories (**Fig 6A**), we discovered a potential functional role that A2 offset responses play in encoding intonation features of meaningful sound categories. Gaining experience caring for mouse pups correlated with weakened A2 On responses elicited by USVs and significantly increased A2 prevalence of USV-evoked Offset Only responses (**Fig 5**), an effect that specifically arose for those USVs with more pup-like frequency trajectories (**Fig 7B**). By testing A_fm_ tuning around natural USVs, we also concluded that only in A2, and only for USVs, did we see a significant overall change from nonmaternal to maternal mice in the tuning to USV frequency trajectory parameters (**Fig 8C**). Although large scale changes in frequency trajectory tuning were not seen here in the Core, plasticity within projection-specific or physiologically distinct subsets of Core neurons, as has been previously reported [32], may occur on a smaller scale but be hidden by population pooling.

Nevertheless, the clear population level difference between Core and A2 after pup care experience suggests that creating a representation of meaningful complex frequency trajectories at the level of A2 must play an important role in interpreting and/or appropriately responding to pup USVs. Interestingly, the shift in A_fm_ tuning we reported results in a larger proportion of maternal A2 neurons with best-A_fm_ near a local peak in the distribution of A_fm_ in pup USVs (Fig 8D). This is reminiscent of operant learning-dependent pure tone BF map expansion seen in Core auditory cortex in many laboratory auditory learning paradigms (e.g. [35, 61]). In our natural learning paradigm, pure tone BF map expansion does not occur in the maternal mouse’s Core auditory cortex [32], perhaps because attention to call onsets, which drives stronger Core BF map plasticity [62], are not as critical for differentiating pup-like from adult-like calls. Instead, there appears to be an “expansion” in the pool of *non-Core* auditory cortical neurons that encodes behaviorally relevant frequency *trajectories*. We also found it intriguing that the location of the maximum slope in A_fm_ tuning curves in the maternal A2 tracked closely to the ideal A_fm_ for discriminating pup-and adult-like USVs based on our logistic regression model. Indeed, the max slope point of a tuning curve theoretically allows the smallest change in A_fm_ to give the largest firing rate change [63].

The fact that representational plasticity in trajectory encoding emerges at the level of A2 could reflect its role as a critical interface between veridical sensory encoding in earlier auditory stages and more perceptually relevant encoding to drive behavioral responses to the naturally variable exemplars of a meaningful sound category. There is some evidence that nonprimary rodent auditory cortex can be more invariant compared to primary auditory cortex to acoustic distortions of natural calls [64]. Primate secondary auditory cortex can also exhibit more categorical responses to trained sound categories [65], and is thought to lie along a hierarchical pathway that produces progressively more categorical responses [66]. By responding to more pup-like frequency trajectories after pup care experience, the maternal A2 may help create a neural representation that is more tolerant of variability in natural call trajectories, so that presumed downstream limbic areas like the amygdala [67, 68] can more categorically drive subcortical circuits for maternal responsiveness [69]. Perhaps in a similar way, intonation differences between emotionally distinct sounds [4, 39] may also be processed through A2 to generate intrinsic or learned physiological responses.

## MATERIALS AND METHODS

### General Methods

Female wildtype CBA/CaJ mice (RRID:IMSR_JAX:000654) between 12-18 weeks of age were used in this study. Animals were socially housed in single-sex cages until breeding age on a 14h light/10h dark reverse light cycle with ad libitum access to food and water; mice were moved to individual housing during experiments. All animal procedures used in this study were approved by the Emory University Institutional Animal Care and Use Committee (IACUC).

At 12-18 weeks of age, mice were moved to individual housing and headpost attachment with small hole craniotomy surgery was conducted [32]. Briefly, animals were anesthetized with isoflurane (2-5%, delivered with oxygen) and buprenorphine (0.1mg/kg) was administered as an analgesic. Animals underwent aseptic surgery to stereotaxically define a recording grid over the left auditory cortex, as the left auditory cortex is putatively associated with mouse vocalization processing, particularly in the maternal paradigm [33, 70]. The skull is exposed, and the left temporal muscle is deflected to permit access to the bone overlying auditory cortex. Using sterile tattoo ink applied to a stiff wire mounted on a stereotaxic manipulator, we marked a grid of ∼100μm diameter dots on the skull in three rows (1.5, 2.0, and 2.5mm below bregma) and five columns (spanning 50-90% of the distance between bregma and lambda, in 10% steps). Dental cement was then used to secure an inverted flat-head screw on the midline equidistant from bregma and lambda. The animal recovered in the home cage placed on a heating pad and was administered saline subcutaneously for fluid replacement.

The day before a recording, the animal was reanesthetized with isoflurane and holes (∼150μm in diameter) were hand drilled on one or more grid points, and a ground hole was drilled over the left frontal cortex. The animal was also acclimated to a foam-lined cylindrical (∼3cm diameter) restraint device, which secured the body while leaving the head exposed. The implanted head post was then secured to a post mounted on a vibration-isolation table, with the restraint device suspended from rubber bands to keep the body in a comfortable position while reducing torque on the headpost. Recordings typically lasted 2-4h, and excessive movement or signs of stress signaled the end of an experiment.

Electrophysiological activity in the auditory cortex was recorded (sample rate 24414.0625/s) in Brainware (TDT) with single 6 MO tungsten electrodes (FHC), filtered between 300Hz and either 3 or 6 kHz. Using a hydraulic Microdrive (FHC), the electrode was driven orthogonally into auditory cortex to an initial depth of ∼200μm, and then advanced in 5um steps until an SU was detected. SU isolation was based on the absence of spikes during the absolute refractory period (1ms), and on online cluster analysis of various spike features (e.g., first vs second peak amplitudes, peak-peak times). In some cases, multiple SUs were recorded at one location and could be extracted by clustering based on spike features.

### Frequency Modulation Tuning around Pure Tone BFs

#### Experimental Details

##### Animal Groups

Besides using intact female mice of the strain and age described in the overall methods, a subset of animals used for sFM tuning around BF were also part of a separate pilot study and had been ovariectomized and given either systemic beta-estradiol or oil vehicle subcutaneous implants. While removing these animals did not change the overall results found, they were included in this section of the analysis for greater statistical power, and to illustrate the ubiquity of neural tuning to short frequency modulations around the BF.

##### Sound Stimulus Playback

Pure tone frequency tuning curves were first obtained with 60 ms duration tone pips at 6 sound levels (15 to 65dB SPL) and 30 frequencies (log-spaced 5-80 kHz), repeated 15 times each and presented in pseudorandom order. To assess A_fm_-only tuning, 60 ms duration sFM stimuli (**Fig 6A**) with f_0_ equal to the BF (f_slope_=0, φ=0) were then played back with 9 steps of increasing A_fm_ (octaves: 0, 1/160, 1/80, 1/40, 1/20, 1/10, 1/5, 1/2, 1). Each sFM sound, presented at the best pure tone level, was repeated 50 times in randomly interleaved trials. To assess combined A_fm_ and f_fm_ tuning, 60 ms duration sFM stimuli with f_0_ equal to the BF (f_slope_=0, φ=0) were presented. f_fm_ was varied in 8 logarithmic steps from 15-137 Hz, while A_fm_ was varied logarithmically from 0-1/4 octave (0, 1/154, 1/82, 1/45, 1/24, 1/13, 1/7, 1/4). A total of 25 randomly interleaved trials per stimulus (presented at the best pure tone level) were collected. For each of these stimuli, we also presented a corresponding bandwidth matched noise stimulus of the same duration, with noise generated in real time with RPvdsEx in Brainware (TDT). The amplitude of the noise was scaled to match the RMS of the corresponding sFM stimulus. A total of 280 pairs of sFM and matched noise stimuli were played to n=40 SUs in n=21 animals. An additional linear FM (lFM) control was used, where an sFM model of a mouse USV was paired with only the linear component of the same sFM model. A total of 35 pairs of sFM and lFM were played to n=20 SUs in n=13 animals.

For all stimuli described, each trial lasted 600 ms, with stimulus playback beginning at 200 ms during a trial.

#### Data Analysis

##### Pure Tone Integration Model for Prediction of A_fm_ Spike Rates

In order to predict spike rates for A_fm_ stimuli (**Fig 1**) from a SU’s pure tone tuning, we computed the power spectral density (MATLAB, pwelch.m) of each of the A_fm_ stimuli. The amount of power in each frequency bin of the power spectrum was multiplied by corresponding frequency bin’s spike rate in the SU’s pure tone tuning curve, normalized such that the evoked response from the BF pure tone for both the A_fm_ tuning stimulus (A_fm_=0) and the pure tone frequency tuning curves were matched. As the number of frequencies played back during pure tone tuning is fewer than the number of (positive) frequency bins in the power spectrum (n=129), the pure tone tuning curve was interpolated linearly across missing frequency bins.

##### Analysis of A_fm_ and f_fm_ Tuning around Best Frequency

We analyzed the 8×8 A_fm_ and f_fm_ tuning stimulus responses to test whether SUs show preference for sFM stimuli over a pure tone. Data used in this analysis included n=61 SUs recorded from n=31 animals. SU responses were divided into the On Resp (during 60 ms window of stimulus playback) and the Offset Resp (window defined starting after stimulus ends until the response drops back to spontaneous levels). n=52 SUs exhibited an On Resp and n=50 exhibited an Offset response. Absolute spike rates were calculated over the On and Offset window for each stimulus. To determine whether a Resp to a specific combination of A_fm_ and f_fm_ significantly differed from that for its BF pure tone, the collection of spike rates from all trials of a given A_fm_ and f_fm_ combination were compared to the collection of spike rates from all pure tone trials using the nonparametric Wilcoxon Rank Sum test. To account for multiple testing, the Bonferroni correction was applied, with the alpha level taken as (0.05/64) or 0.00078. A SU was considered to prefer sFM over pure tone if there was at least one set of sFM parameters (where A_fm_ was non-zero) with significantly greater response than the BF pure tone for either its On or Offset Resp. Comparisons of the number of On vs Offset Resps that had a significantly greater response to at least one sFM stimulus compared to pure tone was conducted via Fisher’s Exact Test.

##### sFM Matched Noise and Linear FM Analysis

For comparison of responses between sFM and matched-bandwidth noise stimuli, responses were calculated during the On, Offset, and the entire evoked response windows, and normalized by subtracting the spontaneous rate, as defined by the spike rate during a 200 ms pre-stimulus silent period. A nonparametric Paired Wilcoxon Signed Rank Test was performed comparing the sFM and paired noise normalized spike rate. In this analysis, a total of n=280 paired sFM and noise stimuli were played to n=40 SUs in n=21 animals. For comparing between sFM and linear FM, we also calculated the responses during the On, Offset, and entire evoked response time windows, normalizing by subtracting the spontaneous rate, followed by performing the Paired Wilcoxon Signed Rank Test. For the linear FM control, a total of n=35 paired sFM and lFM stimuli were played to n=20 SUs from n=13 animals. As a prerequisite to be included for this analysis, all SUs used here showed an excited response to the stimuli, as judged by 2 independent scorers (disagreements, which happened less than 5% of the time for call-excited responses, were not included in the analysis).

### USV Frequency Trajectory Tuning Plasticity

#### Experimental Details

##### Animal Groups

Mice had varying levels of pup experience. The non-maternal animal group consisted of pup-naïve females, pup-naïve females that have only been passively exposed to pup USVs without social interaction, and females that had previously acted as cocarers with a mother, but at the late post-weaning time point (>P21) for electrophysiology, pup calls were no longer salient, based on the lack of preferential phonotaxis to pup USVs [24]. The maternal animal group consisted of post-weaning primiparous mothers (P21, which still find the pup USVs salient [24]), as well as cocarers who were just caring for a litter of pups up to P5-P7 before electrophysiology. For all mice in the maternal group, pup retrieval was conducted on postnatal days P5-P7, in which pups were scattered in the home cage and animals were given 5 minutes to retrieve pups back to the nest. Pup scattering was repeated for a total of three times. Only animals that performed pup retrieval successfully were included in the maternal animal group. No hormonally manipulated animals were included in this section.

##### sFM Modeling of USVs

Mouse pup and adult USVs are complex, single-frequency whistles that have naturally variable frequency trajectories. In order to capture and parameterize the various frequency trajectories of mouse USVs, we used a parameterized sinusoidal plus linear frequency modulated tone model (sFM) to fit to the frequency trajectory of each call (Fig 6A). The parameterized model contains a total of six parameters: duration (dur), onset frequency (f_0_), sFM amplitude (A_fm_), sFM frequency (f_fm_), sFM phase (<p), and linear FM slope (f_slope_). Parameters were fit to minimize the mean squared error between the model and call (reported error = Sqrt(sum squared error / number of samples)). For playback during neural recording, we used a curated natural USV stimulus set containing 18 pup and 18 adult, ground-truth USVs that were matched for duration, onset frequency, and degree of frequency modulation (FM) at onset [32]. In synthesizing the sounds, we applied the original amplitude modulation of the natural USV.

##### Sound Stimulus Playback

USVs (n=36) plus a silent stimulus were played to animals, with up to 50 trials per stimulus, randomly interleaved, as described previously [32]. For a subset of SUs, we also played back the original set of 36 USVs plus 36 sFM models of each USV, randomly interleaved for a total of 25 trials per stimulus.

For each neuron that showed an evoked response to USVs, we then presented additional sFM stimuli optimized around the call that elicited the best response, in order to assess tuning for sFM parameters. This stimulus contained sFM exemplars with f_0_, f_slope_, f_fm_, φ, and dur equal to that of the call eliciting the best response, while A_fm_ was varied in 19 logarithmic steps across the range of natural A_fm_ values in USVs (Afm = 180.78 * exp(0.1051*n); n = [2:2:38]). The original best-response call’s A_fm_ was also included, for a total of 20 different randomly interleaved stimuli in that set, repeated a total of 30 trials per stimulus. If an SU responded to both pup and adult USVs, two A_fm_ tuning stimuli were presented, with one centered around the best pup USV and the other around the best adult USV. In some instances, we also played back a control stimulus set with noise spectrally matched to each of the 20 A_fm_ tuning stimuli. Within the spectrally matched noise control stimulus, for each corresponding A_fm_ tuning stimulus (n=20), 3 instances of randomly generated white noise, each with different random seeding, were first generated and then filtered based on the spectral content of the original A_fm_ stimulus, for a total of 60 stimuli. Each of the 60 stimuli were played for a total of 10 trials per stimulus, such that total presentation time approximately matched that of the sFM-only A_fm_ tuning stimulus. Our ability to hold single units over the course of this protocol varied from site to site, so not all stimuli were played to all units.

#### Data Analysis

##### Spike Density Plot

We generated a spike density plot (MATLAB, dscatter.m, freely available online [71]) to visualize the overall population spiking activity across the n=36 natural mouse USVs, sorted by increasing USV duration (**Fig 4B**). Default smoothing settings were used, in which smoothing was conducted over 20 bins, with the time axis divided into 0.1ms bins (6000 bins across a 600ms period). For visualization purposes, the y axis divided into 100 bins per stimulus (3600 bins for 36 stimuli), and randomized jitter [0-1 bins] was applied to each individual spike along the y axis. A total of n=136 call-excited SUs were included, pooling across region and animal group.

##### On / Offset Prevalence Analysis

In analyzing the prevalence of On and Offset Resps to USVs on a per-call basis (**Fig 5**), the 12 shortest duration calls in the stimulus library were excluded (∼12-15 ms duration calls with faded numbers in the bottom row of Fig 4A), as On and Offset Resps to these short calls could not be definitively differentiated. On and Offset Resps to individual calls were classified by two independent investigators (KC and DK); spiking above spontaneous levels *during* the call’s playback signified an On Resp, while spiking above spontaneous levels *after* the call ended signified an Offset Resp. When both On and Offset Resps were noted for the same call, that response was deemed On+Offset. Data used in this analysis came from n=3264 call responses (n=971 responses were call-excited) played to n=136 call-excited SUs from n=55 animals.

For analyses on a per-SU basis (**Fig 4F**), SUs were classified as On Only, Offset Only or On+Offset SUs based on their call-excited responses to the 24 longer-duration calls. On and Offset Resps of an SU did not necessarily have to be from the same call for that SU to be classified as On+Offset.

##### Logistic Regression Modeling for Classification of sFM Parameter Combinations as Pup-Typical vs Pup-Nontypical

A nominal logistic regression was performed using our library of 10,353 adult and 57,989 pup ultrasonic calls, where calls shorter than 4ms duration and minimum frequencies less than 45kHz were excluded. The nominal logistic regression model was fit using the six sFM parameters (A_fm_, f_fm_, <p, f_0_, f_slope_, dur) to best predict “pup” (pup-like) and “adult” (adult-like) labels. The model followed the format: Score = logit(β_0_ + β_1_ A_fm_ + β_2_ f_fm_ + β_3_ φ + β_4_ f_0_ + β_5_ f_slope_ + β_6_ dur), where each of six beta coefficients were fit to maximize prediction accuracy. A Receiver Operating Characteristics (ROC) curve was constructed using the resulting model, and a threshold Score was selected that maximized the sensitivity and minimized 1-specificity. Significance of the ROC was assessed using a bootstrap analysis (N=1000). All analyses were performed using JMP Pro 13 (SAS).

##### A_fm_ Tuning Analysis

A Gaussian fit 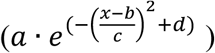 was applied (MATLAB fit.m) to the spike rate as a function of A_fm_ to determine the peak (Gaussian mean, *b*) and bandwidth (Gaussian standard deviation or std, *c*) of a SU’s A_fm_ tuning. Fit parameter initial values and [lower bound, upper bound] were as follows: a = 1 [0,Infinity], b = empirical best A_fm_ (i.e. gives the max spike rate) [0,10000], c = empirical A_fm_ halfmax to halfmax width [0,10000], d = 0 [0,Inf]. Reported best A_fm_ values were taken from the Gaussian fit tuning curves. Note that when using the A_fm_ corresponding with the maximum measured spike rate instead of a Gaussian fit, the results remained the same as reported, but the fit provided a way to estimate the point of maximum slope in the tuning curve (i.e. the standard deviation of the Gaussian).

Results were divided by animal experience group (maternal or nonmaternal) as well as auditory cortical region (Core or A2), and group comparisons were conducted with a nonparametric Wilcoxon Kruskal-Wallis Test followed by the post-hoc Tukey Kramer HSD method, with p<0.05 taken as the significance level. A_fm_ tuning around USV parameters was measured for n=25 calls played to n=18 SUs from n=15 animals. Note when conducting analysis by call, by SU, or by animal, results remained significant.

